# Development of esterase-resistant and highly active ghrelin analogs via thiol-ene click chemistry

**DOI:** 10.1101/2022.06.29.498187

**Authors:** Hao-Zheng Li, Xiao-Xia Shao, Li-Li Shou, Ning Li, Ya-Li Liu, Zeng-Guang Xu, Zhan-Yun Guo

**Author notes:** **Correspondence to Z.-Y. Guo,** Research Center for Translational Medicine at East Hospital, School of Life Sciences and Technology, Tongji University, 1239 Siping Road, Shanghai 200092, China.

## Abstract

The orexigenic peptide ghrelin exerts important functions in energy metabolism and cellular homeostasis by activating the growth hormone secretagogue receptor type 1a (GHSR1a), and thus has therapeutic potential to treat certain diseases. Native ghrelin carries an essential *O*-fatty acyl moiety at the side-chain of its third Ser residue; however, this posttranslational modification is susceptible to hydrolysis by certain esterases in circulation, representing a major route of *in vivo* inactivation of ghrelin. In the present study, we developed a novel approach to prepare various esterase-resistant ghrelin analogs via photo-induced thiol-ene click chemistry. A recombinant unacylated human ghrelin mutant carrying a unique Cys residue at the third position was reacted with commercially available end alkenes, thus various alkyl moieties were introduced to the side-chain of its unique Cys residue via a thioether bond. Among eleven *S*-alkylated ghrelin analogs, analog **11**, generated by reacting with 2-methyl-1-octene, not only acquired much higher stability in human serum and fetal bovine serum, but also acquired moderately higher activity compared with native human ghrelin. Thus, the present study not only provided an efficient approach to prepare various esterase-resistant ghrelin analogs, but also produced a novel highly stable and highly active ghrelin analog with therapeutic potential.

## 1. Introduction

The orexigenic peptide ghrelin is an endogenous agonist of the growth hormone secretagogue receptor type 1a (GHSR1a), an A-class G protein-coupled receptor that was first identified as the receptor of certain synthetic growth hormone secretagogues, hence its name [1,2]. *In vivo*, mature ghrelin is derived from prepropeptide precursors after a series of posttranslational processes, including a special *O*-acylation [1], in which a fatty acyl moiety, typically n-octanoyl, is covalently attached to the side-chain of a Ser residue at the third position via an ester bond (Fig. 1A). This special posttranslational modification is catalyzed by ghrelin *O*-acyltransferase (GOAT), also known as membrane-bound *O*-acyltransferase domain containing 4 (MBOAT4) [3,4]. Recent studies have demonstrated that liver-expressed antimicrobial peptide 2 (LEAP2 or LEAP-2) functions as a competitive antagonist of the ghrelin receptor GHSR1a in mammals and fish [5□11]. The ghrelin system plays important functions in energy metabolism and cellular homeostasis [12‒15], and thus ghrelin and its analogs have therapeutic potential to treat certain diseases, such as anorexia, cancer cachexia, and growth hormone deficiency.

**Fig. 1.**
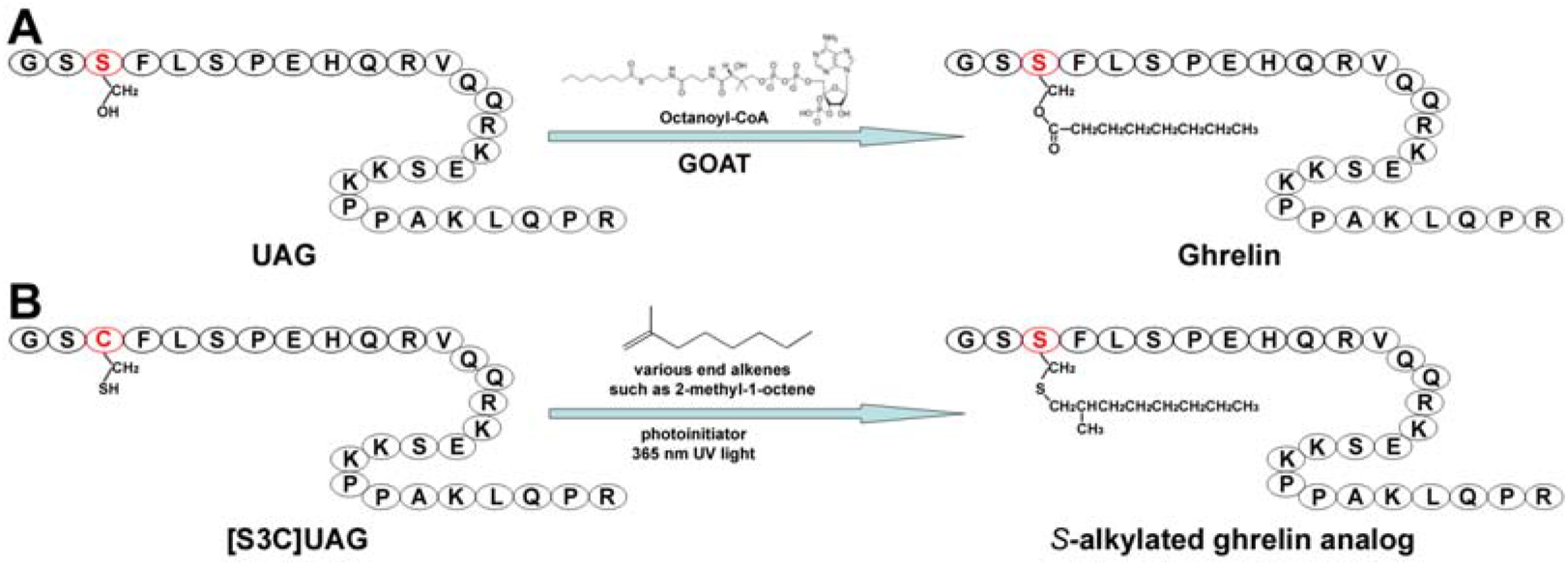
Schematic presentation of *in vivo* octanoylation of UAG by GOAT (**A**) and *in vitro S*-alkylation of [S3C]UAG by the photo-induced thiol-ene click chemistry (**B**). The amino acids in UAG and [S3C]UAG are shown as one-letter code, and the side-chain of their third residue is shown.

Unacylated ghrelin (UAG), also known as des-acyl ghrelin (DAG), has no detectable binding with the ghrelin receptor GHSR1a, although some reports suggested it might have certain biological functions [16□18]. Thus, the special *O*-fatty acyl modification is essential for ghrelin binding to, and activation of, its receptor GHSR1a [1]. However, the *O*-fatty acyl moiety of ghrelin is susceptible to hydrolysis by certain esterases in circulation [19–22], such as butyryl cholinesterase, carboxylesterase, *α*_2_ macroglobulin, and acyl-protein thioesterase 1 (also known as lysophospholipase 1). Hydrolysis of the fatty acyl moiety represents a major route of *in vivo* inactivation of ghrelin; therefore, the development of esterase-resistant ghrelin analogs with high activity is needed for the therapeutic application of ghrelin.

In a previous study, a chemically synthesized ghrelin analog carrying an n-octyl moiety via a thioether bond displayed only moderately lower activity compared with native ghrelin [23], implying that *S*-alkylation might be a suitable approach to develop esterase-resistant ghrelin analogs. To conveniently prepare various *S*-alkylated ghrelin analogs, in the present study, we employed photo-induced thiol-ene click chemistry, which has been used in peptide modifications in recent years [24–26]. For this purpose, we designed an analog of human UAG, designated as [S3C]UAG, by replacing Ser3 of human UAG with a Cys residue. The mature [S3C]UAG peptide was prepared by overexpression of a large precursor in *Escherichia coli* and subsequent chemical cleavage using cyanogen bromide (Fig. S1 and S2). Thereafter, it was reacted with some commercially available end alkenes and different alkyl moieties were introduced to the side-chain of its unique Cys residue via a thioether bond (Fig. 1B). Using this approach, we generated eleven *S*-alkylated ghrelin analogs in the present study. Among them, analog **11** not only acquired markedly higher stability in human serum and fetal bovine serum, but also acquired moderately higher activity compared with that of native human ghrelin. Thus, the present study not only provided an efficient approach to prepare various esterase-resistant ghrelin analogs, but also developed a novel highly stable and highly active ghrelin analog with therapeutic potential.

## 2. Materials and methods

### 2.1. Chemical synthesis of native human ghrelin

The native human ghrelin was chemically synthesized at GL Biotech (Shanghai, China) via solid-phase peptide synthesis using standard Fmoc methodology. The crude peptide was purified to homogeneity via high performance liquid chromatography (HPLC) sequentially using a semi-preparative C_18_ reverse-phase column (Zorbax 300SB-C18, 9.4 × 250 mm; Agilent Technologies, Santa Clara, CA, USA) and an analytical C_18_ reverse-phase column (Zorbax 300SB-C18, 4.6 × 250 mm; Agilent Technologies). The synthetic ghrelin was eluted from the reverse-phase columns by an acetonitrile gradient composed of solvent A (0.1% aqueous TFA) and solvent B (acetonitrile containing 0.1% TFA). Identity of the synthetic native human ghrelin was confirmed by electrospray mass spectrometry.

### 2.2. Preparation of the recombinant [S3C]UAG

The nucleotide sequence encoding the fusion protein of [S3C]UAG and human C4ORF48 peptide was chemically synthesized at GeneWiz (Suzhou, China). After cleavage with restriction enzymes NdeI and EcoRI, the synthetic DNA fragment was ligated into a pET vector, resulting in the construct pET/6×His-C4ORF48-[S3C]UAG. The nucleotide sequence and amino acid sequence of the fusion protein are shown in supplementary Fig. S1.

For bacterial overexpression, the plasmid construct was transformed into *E. coli* strain BL21(DE3), and the transformed bacteria were cultured in liquid Luria-Bertani medium (with 100 *μ*g/ml ampicillin) to OD600 ≈ 2.0 at 37°C with vigorous shaking. Thereafter, isopropyl β-D-thiogalactoside was added to the final concentration of 1.0 mM, and the bacteria were continuously cultured at 37°C for 5−6 h with gentle shaking. Subsequently, the *E. coli* cells were harvested by centrifugation (5000 *g*, 10 min), re-suspended in lysis buffer (20 mM phosphate buffer, pH 7.4, 0.5 M NaCl), and lysed by sonication. Thereafter, inclusion bodies were collected by centrifugation (18000 *g*, 30 min), re-suspended in solubilizing buffer (lysis buffer plus 6 M guanidine chloride), and subjected to *S*-sulfonation by adding solid sodium sulfite (Na_2_SO_3_) and sodium tetrathionate (Na_2_S_4_O_6_) to the final concentrations of 100 mM and 80 mM, respectively. After shaking at room temperature for ∼3 h, the supernatant was applied to an immobilized metal ion affinity chromatography, and the *S*-sulfonated fusion protein was eluted from a Ni^2+^ column by 250 mM imidazole (in solubilizing buffer). The eluted fraction was then dialyzed against water overnight, and precipitate of the fusion protein was collected by centrifugation (8000 *g*, 10 min).

Thereafter, the *S*-sulfonated fusion protein was dissolved in cleavage solution (0.1 M hydrochloride, 2 M guanidine hydrochloride) at the concentration of ∼10 mg/ml, and mixed with equal volume of ∼10 mg/ml CNBr solution (dissolved in cleavage solution). After incubation at room temperature for ∼15 min, the reaction mixture was 10-fold diluted with 20% acetronitrile, its pH was adjusted to 3−4 by adding appropriate amount of 1.0 M NaOH solution, and then applied to HPLC. Peptide fractions were eluted from a semi-preparative C_18_ reverse-phase column (Zorbax 300SB-C18, 9.4 × 250 mm; Agilent Technologies) by an acetonitrile gradient. The eluted fractions were manually collected, lyophilized, and their identities were confirmed by electrospray mass spectrometry.

The lyophilized *S*-sulfonated [S3C]UAG fraction was dissolved in 100 mM Tris-Cl buffer (pH8.5) at the final concentration of ∼2 mg/ml, and treated with 50 mM dithiothreitol at room temperature for ∼30 min. Thereafter, pH of the reaction mixture was adjusted to 3−4 by adding appropriate amount of TFA, and then applied to HPLC. The mature [S3C]UAG was eluted from an analytical C_18_ reverse-phase column (Zorbax 300SB-C18, 4.6 × 250 mm; Agilent Technologies) by an acetonitrile gradient, manually collected, lyophilized, and confirmed by electrospray mass spectrometry.

### 2.3. S-alkylation of [S3C]UAG via photo-induced thiol-ene click chemistry

All photo-induced thiol-ene reactions were conducted in an Aldrich^®^ AtmosBag (Sigma-Aldrich, Saint Louis, MO, USA) filled with nitrogen gas. A 365 nm ultra-violet light emitting diode (UV-LED) lamp was used to initiate the thiol-ene reaction, which was conducted according to previously published procedures [27,28], using the hydrophilic 2-hydroxy-4’-(2-hydroxyethoxy)-2-methylpropiophenone (CAS no. 106797-53-9) as a photo initiator. The reaction components, including lyophilized [S3C]UAG, -enes, the photo initiator, triisopropylsilane (CAS no. 6485-79-6), and tert-dodecylmercaptan (CAS no. 25103-58-6) were all dissolved in 1-methyl-2-pyrrolidinone (CAS no. 872-50-4) as stock solutions. To start the reaction, these components were mixed together in a small open glass bottle at the following concentrations: [S3C]UAG, 1.9 mM; -ene, 19 mM; the photo initiator, 9 mM; triisopropylsilane, 47 mM, and tert-dodecylmercaptan, 25 mM. The final reaction solvent contained 80% (v/v) 1-methyl-2-pyrrolidinone, 15% (v/v) water, and 5% (v/v) TFA. After addition of these components, the glass bottle was put into the AtmosBag, which was then filled with nitrogen gas. Thereafter, the bottle was irradiated using the UV-LED lamp (∼60 mW/cm^2^) in the AtmosBag at room temperature for 45 min or for the indicated times. After the UV light exposure, the reaction mixture was 10-fold diluted with 20% acetonitrile, its pH was adjusted to 3−4 by adding appropriate amount of 1.0 M NaOH solution, and it was then applied to HPLC. Fractions eluted from an analytical C_18_ reverse-phase column (Zorbax 300SB-C18, 4.6 × 250 mm; Agilent Technologies) using an acetonitrile gradient were manually collected, lyophilized, and their identities were confirmed by electrospray mass spectrometry.

### 2.4. Preparation of tracers for receptor-binding assays

The SmBiT-based tracers, ghrelin-SmBiT and LEAP2-SmBiT, for the NanoBiT-based homogenous binding assays were prepared according to our previous procedure by chemical conjugation of a C-terminally Cys-tagged SmBiT with a C-terminally Cys-tagged human ghrelin or a C-terminally Cys-tagged human LEAP2 mutant via an intermolecular disulfide linkage [6,29]. The NanoLuc-conjugated ghrelin tracer (ghrelin-Luc) for the washing-based binding assay was prepared according to our previous procedure by chemical conjugation of a C-terminally Cys-tagged NanoLuc luciferase with a C-terminally Cys-tagged human ghrelin via an intermolecular disulfide linkage [30].

### 2.5. Receptor-binding assays of the S-alkylated ghrelin analogs

Binding activity of the *S*-alkylated ghrelin analogs with human GHSR1a was measured using the NanoBiT-based homogenous binding assay according to our previous procedures [29,31]. Briefly, human embryonic kidney (HEK) 293T cells were transiently transfected with the expression construct pcDNA6/sLgBiT-GHSR1a that encodes an N-terminally secretory large NanoLuc fragment (sLgBiT)-fused human GHSR1a. Next day, the transfected cells were trypsinized, seeded into white opaque 96-well plates, and cultured for ∼24 h to ∼90% confluence. To conduct the homogenous binding assay, medium was removed and binding solution for the NanoBiT-based binding assay (serum-free DMEM plus 0.1% BSA and 0.01% Tween-20) was added (50 *μ*l/well). The binding solution contained a constant concentration of tracer (ghrelin-SmBiT or LEAP2-SmBiT) and varied concentrations of native human ghrelin or the *S*-alkylated analogs. After incubation at 24°C for ∼1 h, diluted NanoLuc substrate (Promega, Madison, WI, USA, 30-fold dilution using the binding solution) was added (10 *μ*l/well) and bioluminescence was immediately measured on a SpectraMax iD3 microplate reader (Molecular Devices, Sunnyvale, CA, USA). The measured bioluminescence data were expressed as mean ± standard deviation (SD, *n* = 3) and fitted to one-site binding model using SigmaPlot 10.0 software (SYSTAT software, Chicago, IL, USA).

### 2.6. Receptor activation assays of the S-alkylated ghrelin analogs

Activation potency of the *S*-alkylated ghrelin analogs towards human GHSR1a was measured using a cAMP-response element (CRE)-controlled NanoLuc reporter according to our previous procedures [29,31]. Briefly, HEK293T cells were transiently cotransfected with an expression construct of human GHSR1a (pcDNA6/GHSR1a) and a CRE-controlled NanoLuc reporter vector (pNL1.2/CRE). Next day of the transfection, the cells were trypsinized, seeded into white opaque 96-well plates, and cultured for ∼24 h to ∼80% confluence. To conduct the activation assay, medium was removed and activation solution (serum-free DMEM plus 1% BSA) was added (50 *μ*l/well). The activation solution contained varied concentrations of native human ghrelin or the *S*-alkylated analogs. After the cells were continuously cultured at 37°C for ∼4 h, diluted NanoLuc substrate (Promega, 30-fold dilution using the activation solution) was added (10 *μ*l/well) and bioluminescence was immediately measured on a SpectraMax iD3 microplate reader (Molecular Devices). The measured bioluminescence data were expressed as mean ± SD (*n* = 3) and fitted to sigmoidal curves using SigmaPlot 10.0 software (SYSTAT software).

### 2.7. Stability measurement of native human ghrelin and the most active analog

To measure their stability, native human ghrelin and the most active analog, **11**, were diluted in binding solution (PBS plus 1% BSA) to concentrations of 200, 400, and 800 nM, respectively. Subsequently, the diluted samples were mixed with human serum or fetal bovine serum at a volume ratio of 1:9, and then incubated at 37°C for 15 h. For the standard binding curves, the freshly diluted peptides were mixed with the pre-incubated serum (37°C for 15 h) at a volume ratio of 1:9 just before the binding assay. Thereafter, the serum-mixed peptide samples were mixed with an equal volume of 0.5 nM ghrelin-Luc tracer (diluted in PBS plus 1% BSA), and then added to living HEK293T cells transiently overexpressing human GHSR1a (50 *μ*l/well in 96-well white opaque plates). After incubation at 24°C for ∼1 h, the peptide solutions were removed and the cells were washed twice with ice-cold PBS (200 *μ*l/well/time). Thereafter, diluted NanoLuc substrate (Promega, 100-fold dilution using PBS) was added (50 *μ*l/well) and bioluminescence was immediately measured on a SpectraMax iD3 microplate reader (Molecular Devices). The measured bioluminescence data were expressed as the mean ± SD (*n* = 3) and the standard binding curves were fitted to one-site binding models using SigmaPlot 10.0 software (SYSTAT software). The concentration of the active native human ghrelin or analog **11** remaining after overnight incubation in serum was calculated from their measured bioluminescence according to the standard binding curves.

## 3. Results

### 3.1. Preparation of the S-alkylated ghrelin analogs via the photo-induced thiol-ene click chemistry

To prepare the small [S3C]UAG (28 amino acids) via bacterial overexpression, we designed a larger precursor by fusing [S3C]UAG to the C-terminus of the expected mature peptide (61 amino acids) of human C4ORF48 via a Met residue (Fig. S2A). As expected, the larger precursor could be efficiently overexpressed in *E. coli* as inclusion bodies. To obtain the mature [S3C]UAG peptide from the recombinant precursor, we developed an efficient procedure to purify and process the precursor (Fig. S2B). After the inclusion bodies were solubilized via an *S*-sulfonation approach, the *S*-sulfonated precursor was purified using an immobilized metal ion affinity chromatography and then subjected to CNBr cleavage. As monitored by HPLC (Fig. S2C), the *S*-sulfonated [S3C]UAG could be quickly released from the precursor by chemical cleavage. Finally, the reversibly modified *S*-sulfonate moiety was removed via dithiothreitol treatment, and mature [S3C]UAG was obtained at a considerable yield. From one liter of the *E. coli* culture broth, typically 5−10 mg of mature [S3C]UAG peptide could be obtained within a week.

To introduce various alkyl moieties into the mature [S3C]UAG, we optimized the reaction conditions of the photo-induced thiol-ene click chemistry. We selected the hydrophilic 2-hydroxy-4’-(2-hydroxyethoxy)-2-methylpropiophenone as the photo initiator, because its elution peaks were well separated from those of the *S*-alkylated ghrelin analogs on HPLC (Fig. 2). To initiate the photo reaction, we used a 365 nm UV-LED lamp, which has a sharp spectrum around 365 nm and an adjustable power output. To minimize the oxidized byproducts, we conducted all photo-induced reactions in an AtmosBag filled with nitrogen gas.

**Fig. 2.**
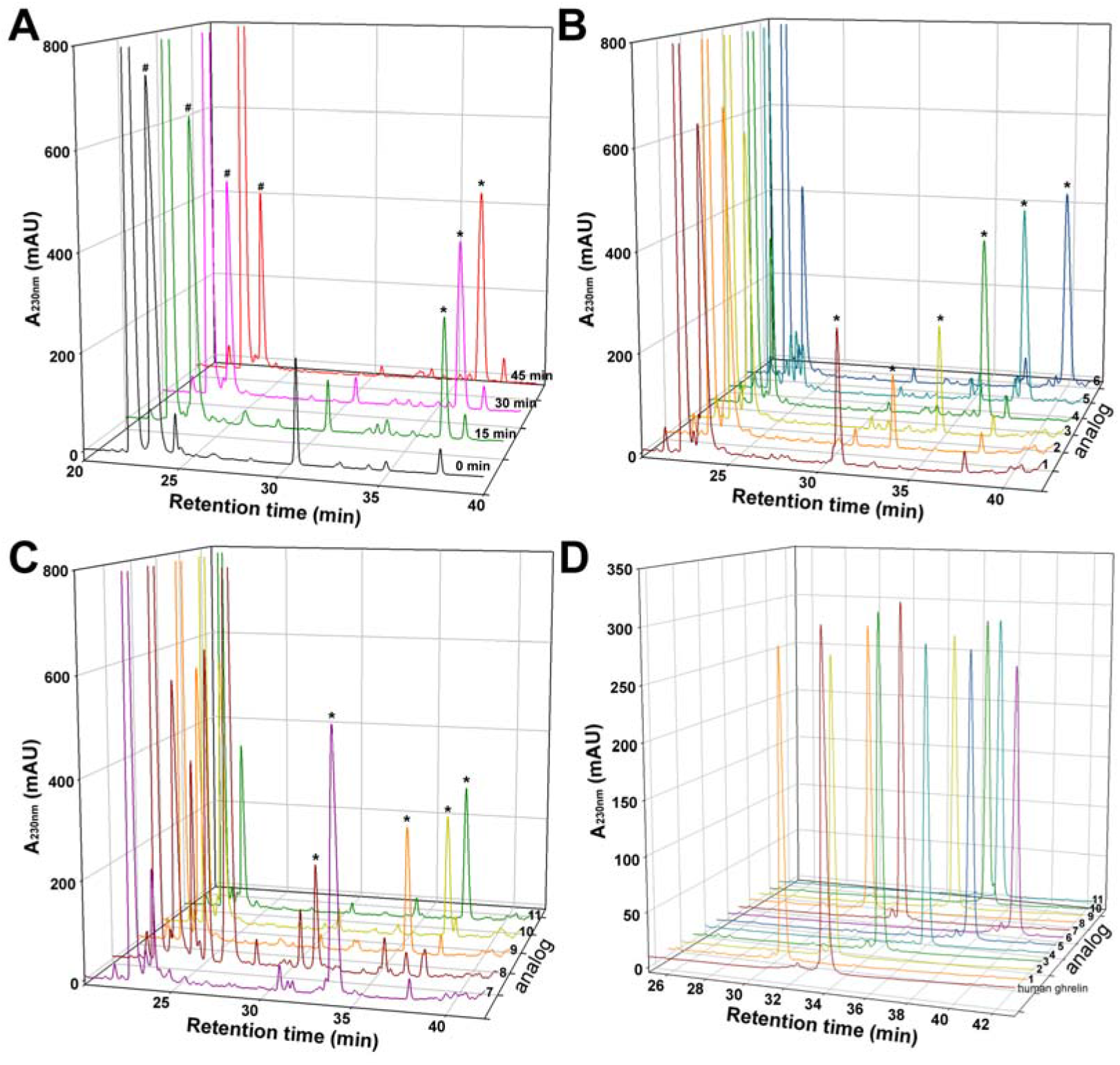
Preparation of the *S*-alkylated ghrelin analogs by photo-induced thiol-ene click chemistry. (**A**) HPLC analyses of the reaction mixture of [S3C]UAG reacting with 1-octene after UV light exposure for different times. The peak of [S3C]UAG is indicated by an octothorpe, and that of the *S*-octylated product is indicated by an asterisk. (**B**,**C**) HPLC analyses of the reaction mixture of [S3C]UAG reacting with various alkenes after UV light exposure for 45 min. The peak of [S3C]UAG is indicated by an octothorpe, and that of the *S*-alkylated analogs is indicated by an asterisk. (**D**) Purity analyses of the *S*-alkylated ghrelin analogs by HPLC.

When [S3C]UAG was reacted with 1-octene, its elution peak (indicated by an octothorpe) on HPLC decreased quickly after exposure to UV light, meanwhile a new peak (indicated by an asterisk) appeared and increased correspondingly (Fig. 2A). As measured by mass spectrometry (Table 1), the new peak was the expected *S*-octylated ghrelin analog. Estimated from their elution peaks, approximately 60% of [S3C]UAG could be converted to the *S*-octylated product (Fig. 2A).

**Table 1.**
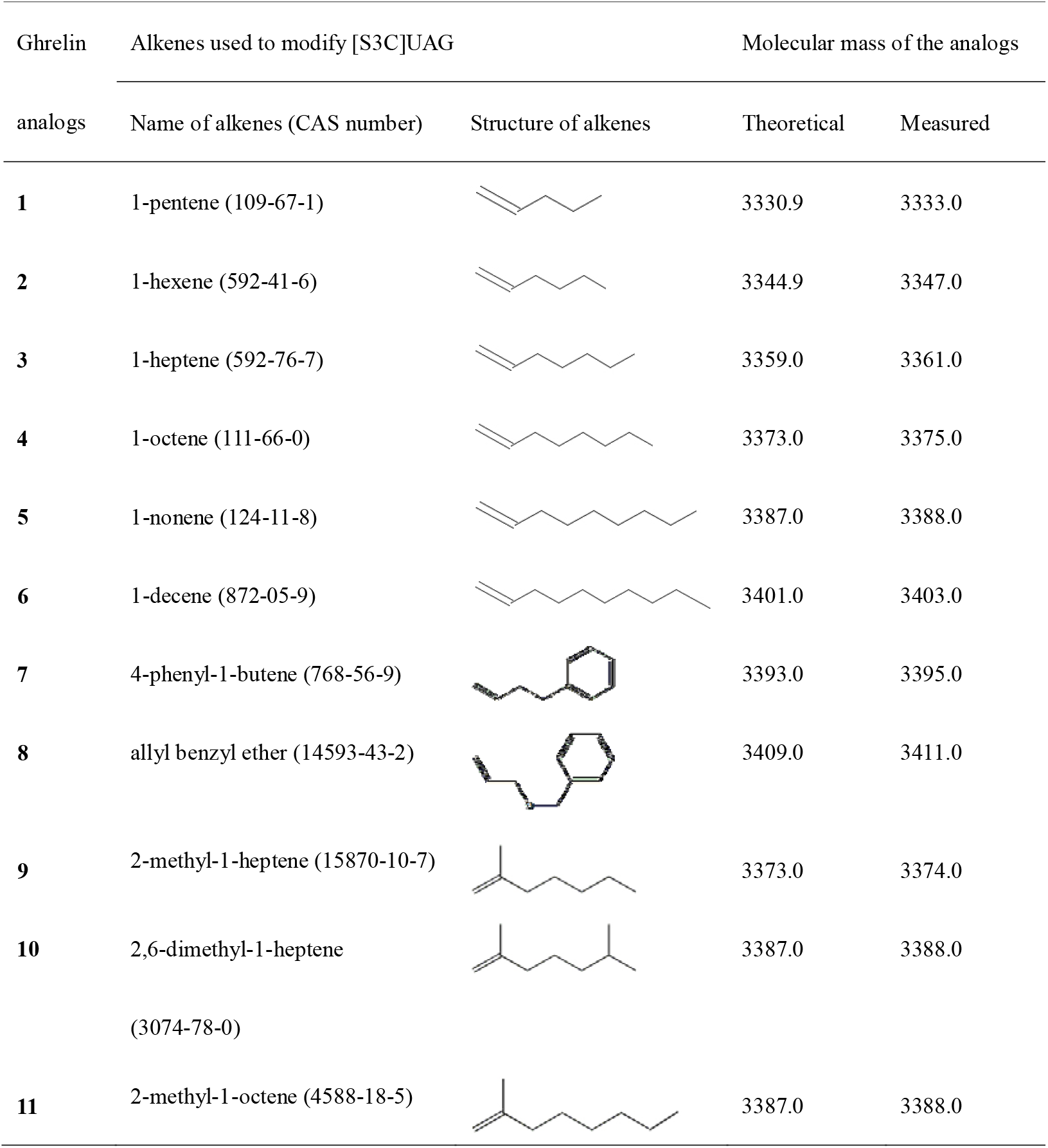
Summary of the measured molecular mass of the *S*-alkylated ghrelin analogs.

| Ghrelin<br>analogs | Alkenes used to modify [S3C]UAG |  | Molecular mass of the analogs |  |
| --- | --- | --- | --- | --- |
|  | Name of alkenes (CAS number) | Structure of alkenes | Theoretical | Measured |
| **1** | 1-pentene (109-67-1) |  | 3330.9 | 3333.0 |
| **2** | 1-hexene (592-41-6) |  | 3344.9 | 3347.0 |
| **3** | 1-heptene (592-76-7) |  | 3359.0 | 3361.0 |
| **4** | 1-octene (111-66-0) |  | 3373.0 | 3375.0 |
| **5** | 1-nonene (124-11-8) |  | 3387.0 | 3388.0 |
| **6** | 1-decene (872-05-9) |  | 3401.0 | 3403.0 |
| **7** | 4-phenyl-1-butene (768-56-9) |  | 3393.0 | 3395.0 |
| **8** | allyl benzyl ether (14593-43-2) |  | 3409.0 | 3411.0 |
| **9** | 2-methyl-1-heptene (15870-10-7) |  | 3373.0 | 3374.0 |
| **10** | 2,6-dimethyl-1-heptene<br>(3074-78-0) |  | 3387.0 | 3388.0 |
| **11** | 2-methyl-1-octene (4588-18-5) |  | 3387.0 | 3388.0 |

When [S3C]UAG was reacted with other end alkenes (Table 1), *S*-alkylated ghrelin analogs were obtained with the yields ranging from ∼30 to ∼80% (Fig. 2B,C). We prepared eleven *S*-alkylated ghrelin analogs in total via the thiol-ene click chemistry, which all displayed the expected molecular mass as analyzed by mass spectrometry (Table 1) and a symmetrical elution peak as analyzed by HPLC (Fig. 2D). Thus, it seemed that various *S*-alkylated ghrelin analogs could be conveniently prepared using the present approach.

### 3.2. Activity of the S-alkylated ghrelin analogs

To measure activity of these *S*-alkylated ghrelin analogs, we employed a receptor binding assay and a receptor activation assay (Fig. 3). We measured their binding activity with human GHSR1a using the NanoBiT-based homogenous binding assay [6,11,29,31], and measured their activation potency towards human GHSR1a using a CRE-controlled NanoLuc reporter [6,11,29,31]. Although GHSR1a signals through the Gq pathway, it also activates the transcription factor CREB through kinases such as Ca^2+^/calmodulin kinase IV and protein kinase C [32]. Thus, a CRE-controlled luciferase reporter can be used to monitor GHSR1a activation, as demonstrated in previous studies [6,31,33].

**Fig. 3.**
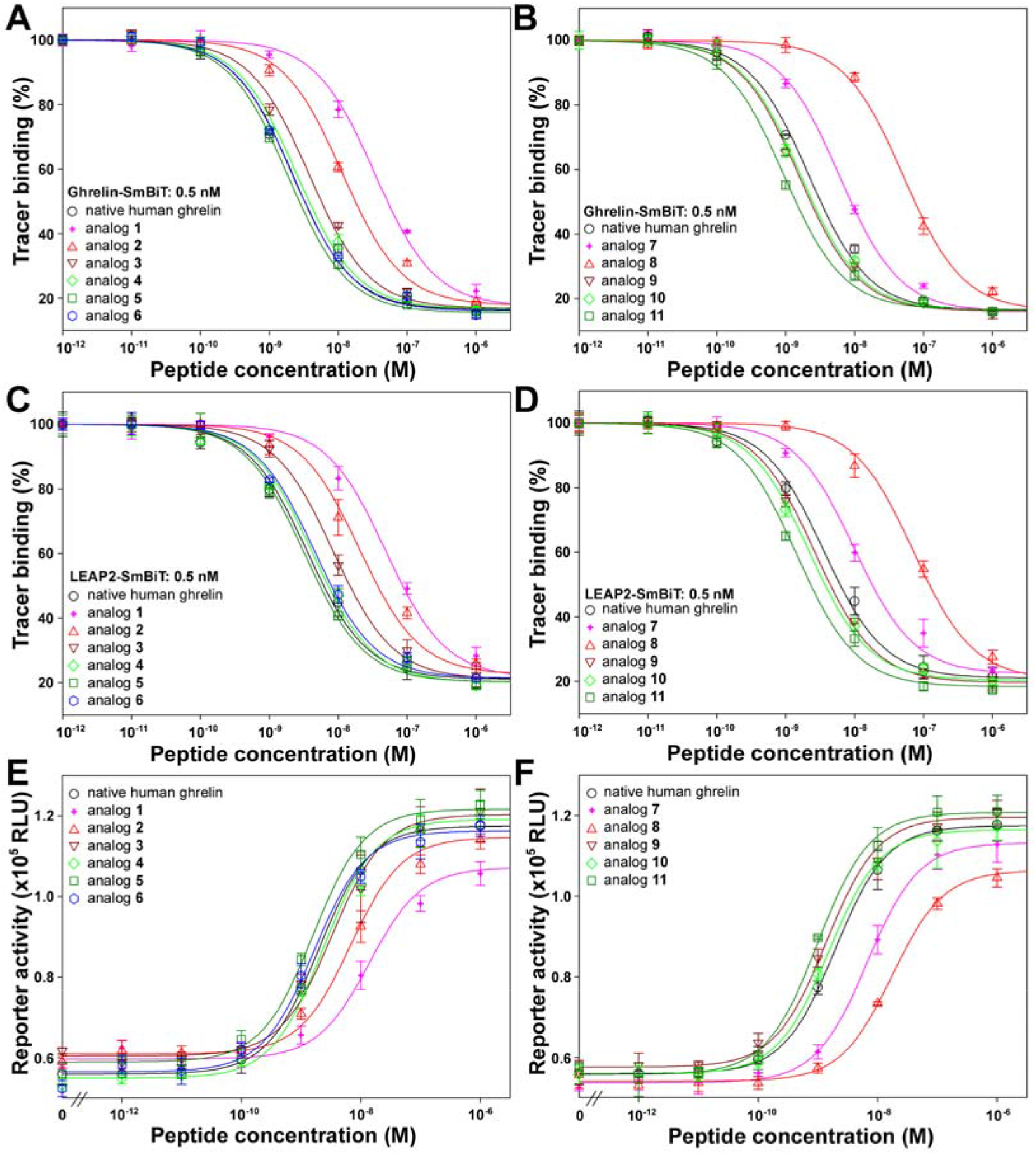
Activity assays of the *S*-alkylated ghrelin analogs. (**A**,**B**) Binding activity with human GHSR1a measured by the NanoBiT-based binding assay using ghrelin-SmBiT as a tracer. (**C**,**D**) Binding activity with human GHSR1a measured by the NanoBiT-based binding assay using LEAP2-SmBiT as a tracer. (**E**,**F**) Activation potency towards human GHSR1a measured by a CRE-controlled NanoLuc reporter.

Compared to native human ghrelin, analog **1** carrying a short n-pentyl moiety displayed ∼15-fold lower receptor binding activity (Fig. 3A,C and Table 2) and ∼7-fold lower receptor activation potency (Fig. 3E and Table 2). When the chain length of the linear alkyl moiety was increased from n-pentyl (-C_5_H_11_) to n-nonyl (-C_9_H_19_), both the receptor binding activity and the receptor activation potency of analogs **2**−**5** increased gradually, with analog **5** carrying an n-nonyl moiety having full activity compared with native human ghrelin (Fig. 3A,C,E and Table 2). However, introduction of a longer n-decyl (-C_10_H_21_) moiety caused slight decrease in activity (Fig. 3A,C,E and Table 2), suggesting that further increase of the chain length of the linear alkyl moiety likely has detrimental effects.

**Table 2.** Summary of the measured IC_50_ and EC_50_ values of the *S*-alkylated ghrelin analogs.

| Ghrelin analogs | IC<sub>50</sub> (nM) |  | EC<sub>50</sub> (nM) |
| --- | --- | --- | --- |
|  | Ghrelin-SmBiT | LEAP2-SmBiT | NanoLuc reporter |
| Native human ghrelin | 2.19 ± 0.15 | 3.55 ± 0.34 | 1.95 ± 0.21 |
| **1** | 33.9 ± 2.3 | 49.0 ± 4.3 | 13.8 ± 1.8 |
| **2** | 11.5 ± 0.8 | 20.4 ± 2.5 | 6.76 ± 0.83 |
| **3** | 3.90 ± 0.25 | 8.51 ± 0.57 | 3.31 ± 0.58 |
| **4** | 2.51 ± 0.17 | 4.27 ± 0.41 | 2.40 ± 0.31 |
| **5** | 1.91 ± 0.09 | 3.24 ± 0.22 | 1.58 ± 0.17 |
| **6** | 2.24 ± 0.15 | 4.57 ± 0.33 | 1.62 ± 0.24 |
| **7** | 6.03 ± 0.28 | 10.0 ± 0.9 | 6.61 ± 0.80 |
| **8** | 50.1 ± 3.3 | 70.8 ± 4.7 | 17.4 ± 1.6 |
| **9** | 1.58 ± 0.10 | 2.69 ± 0.18 | 1.38 ± 0.17 |
| **10** | 1.66 ± 0.11 | 2.19 ± 0.15 | 1.51 ± 0.16 |
| **11** | 0.93 ± 0.07 | 1.45 ± 0.10 | 1.00 ± 0.11 |

When an aromatic phenyl moiety was introduced, the resultant analogs **7** and **8** displayed significantly lower activity compared with native human ghrelin (Fig. 3B,D,F and Table 2), suggesting that the aromatic ring cannot be well accommodated in the ligand-binding pocket of GHSR1a. Analogs **9** and **11**, carrying a branched alkyl moiety with a methyl group at the second carbon atom, displayed significantly higher activity than the corresponding analogs **3** and **4** carrying a linear alkyl moiety (Fig. 3B,D,F and Table 2), suggesting that the methyl branch at the second carbon favors interaction of the analogs with receptor GHSR1a. However, introduction of a branch at the far end of the hydrocarbon chain seemingly had no effects, since analog **9** and **10** displayed similar activity (Fig. 3B,D,F and Table 2). Among these *S*-alkylated ghrelin analogs, analog **11** was most active in both the receptor binding assay and the receptor activation assay, displaying approximately two-fold higher activity than native human ghrelin.

### 3.3. Stability of the most active S-alkylated ghrelin analog

As shown above, analog **11** acquired moderately higher activity compared with native human ghrelin. Next, we wanted to know whether its stability was significantly increased, as expected. To compare its stability with that of native human ghrelin, we incubated both peptides in human serum or fetal bovine serum, and then quantified the remaining active peptide via a washing-based ligand−receptor binding assay using NanoLuc-conjugated ghrelin as a bioluminescent tracer [30]. In the presence of 45% human serum or fetal bovine serum in the binding assay, the standard competition binding curves were typically sigmoidal for both peptides (Fig. 4A,B), suggesting that this washing-based binding assay is resistant to interference from a high percentage of serum. In the presence of human serum or fetal bovine serum, analog **11** and native human ghrelin displayed similar IC_50_ values (Table 3), confirming that the analog is highly active.

**Fig. 4.**
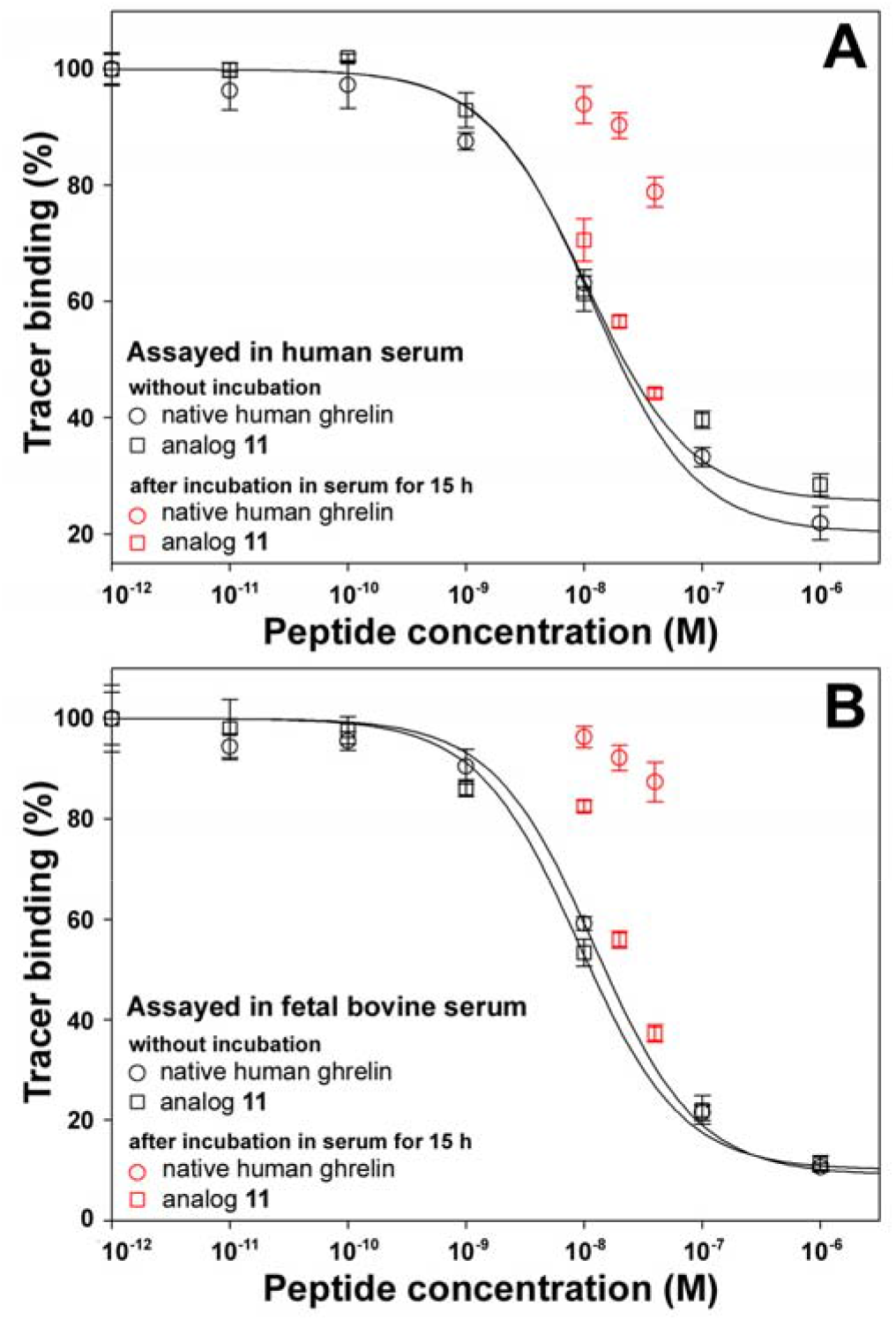
Stability assay of native human ghrelin and analog **11** in human serum (**A**) or fetal bovine serum (**B**) using the washing-based receptor binding assay.

**Table 3.**
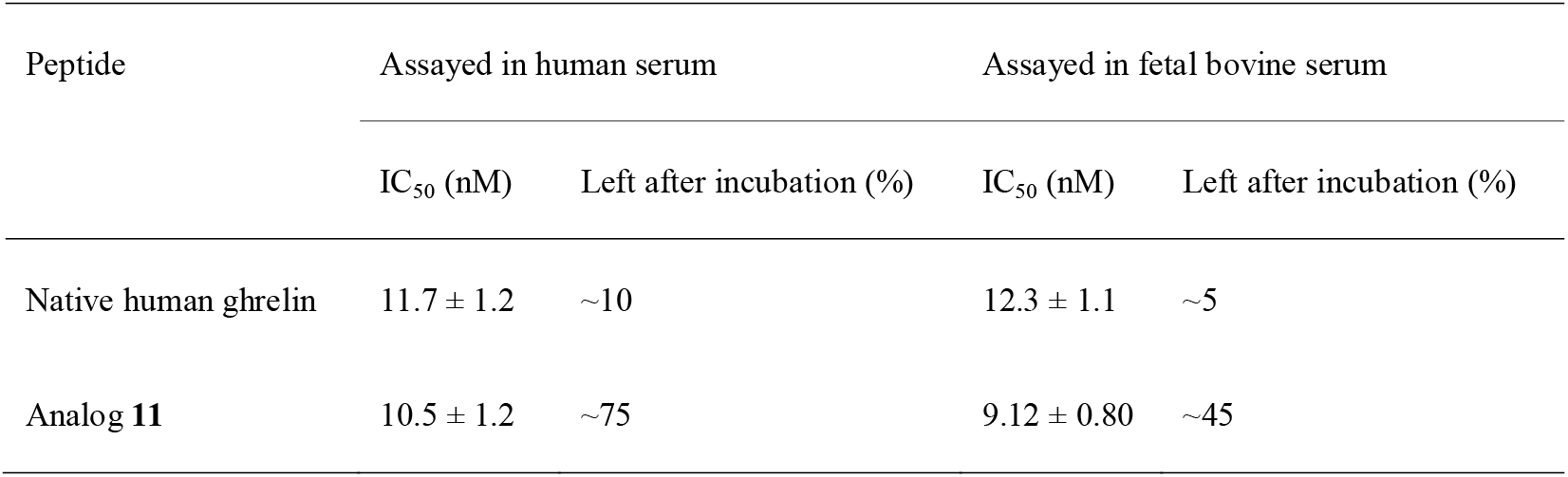
Summary of stability of native human ghrelin and analog **11** in human serum and fetal bovine serum measured by the washing-based receptor binding assay.

| Peptide | Assayed in human serum |  | Assayed in fetal bovine serum |  |
| --- | --- | --- | --- | --- |
|  | IC<sub>50</sub> (nM) | Left after incubation (%) | IC<sub>50</sub> (nM) | Left after incubation (%) |
| Native human ghrelin | 11.7 ± 1.2 | ~10 | 12.3 ± 1.1 | ~5 |
| Analog **11** | 10.5 ± 1.2 | ~75 | 9.12 ± 0.80 | ~45 |

After native human ghrelin was incubated in 90% human serum at 37°C for 15 h, its measured binding values were far from the standard binding curve of the native human ghrelin (Fig. 4A), suggesting that incubation with human serum caused significant inactivation of native human ghrelin. Calculated according to the standard binding curve, only ∼10% of human ghrelin remained active after incubation with 90% human serum at 37°C for 15 h (Table 3). In contrast, incubation with human serum had much less effect on analog **11** (Fig. 4A): ∼70% of the analog remained active after incubation with 90% human serum at 37°C for 15 h (Table 3). A similar phenomenon was observed for both peptides after incubation with fetal bovine serum (Fig. 4B and Table 3). Thus, analog **11** acquired much higher stability in both human serum and fetal bovine serum compared with native human ghrelin, suggesting that the *S*-alkylated analog would have a much longer *in vivo* half-life than native ghrelin when used as a therapeutic reagent in future studies.

## Discussion

In the present study, we developed an efficient approach to prepare highly stable and highly active ghrelin analogs by *S*-alkylation of [S3C]UAG with various alkenes via the photo-induced thiol-ene click chemistry. In future studies, more *S*-alkylated ghrelin analogs could be prepared via this approach. To further improve the activity and stability of the *S*-alkylated analogs, two approaches might be used in future studies. On the one hand, some mutations might be introduced to other positions of the [S3C]UAG peptide to make the peptide chain more resistant to proteases in circulation. Thereafter, the designed analogs might be prepared either by bacterial overexpression or by solid-phase peptide synthesis. On the other hand, more end alkenes with various structures might be used to modify [S3C]UAG or its analogs. Thus, our present study provided a practical approach to prepare various *S*-alkylated ghrelin analogs with high activity and high stability.

In the present study, we generated eleven *S*-alkylated ghrelin analogs via modification of [S3C]UAG with different alkenes. Among them, analog **11**, generated by reacting with 2-methyl-1-octene, displayed the highest activity: its activity was approximately two-fold higher than that of native human ghrelin. Thus, it seemed that the branched methyl group at the second position of the modified alkyl moiety favors the activity of the *S*-alkylated analog. In future studies, the effect of a larger branched group, such as an ethyl group, might be tested using the corresponding alkenes.

After an end alkene with a branch at the second position, such as 2-methyl-1-octene and 2-methyl-1-heptene, was reacted with [S3C]UAG, its second carbon will be converted to a chiral atom with an R-configuration or an S-configuration. Thus, the reaction theoretically produces two *S*-alkylated analogs carrying an alkyl moiety either with an R-configuration or with an S-configuration at the second carbon atom. However, only one elution peak was identified as the modification product when [S3C]UAG was reacted with these branched alkenes, suggesting that the two expected products could not be separated on HPLC. Thus, it is unclear whether the two products have similar activity or not. In future studies, the corresponding analogs might be chemically synthesized in order to test their activity.

## Acknowledgments

This work was supported by grant from the National Natural Science Foundation of China (31971193).

## Abbreviations

BSA: bovine serum albumin
CRE: cAMP-response element
DAG: des-acyl ghrelin
GHSR1a: growth hormone secretagogue receptor type 1a
DMEM: Dulbecco’s Modified Eagle Medium
GOAT: ghrelin *O*-acyltransferase
HEK: human embryonic kidney
HPLC: high performance liquid chromatography
LEAP2: liver-expressed antimicrobial peptide 2
LED: light emitting diode
MBOAT4: membrane-bound *O*-acyltransferase domain containing 4
NanoBiT: NanoLuc Binary Technology
PBS: phosphate-buffered saline
SD: standard deviation
sLgBiT: secretory large NanoLuc fragment for NanoBiT
SmBiT: low-affinity complementation tag for NanoBiT
TFA: trifluoroacetic acid
UAG: unacylated ghrelin
UV: ultra-violet

## Supplementary materials

**Fig. S1.**
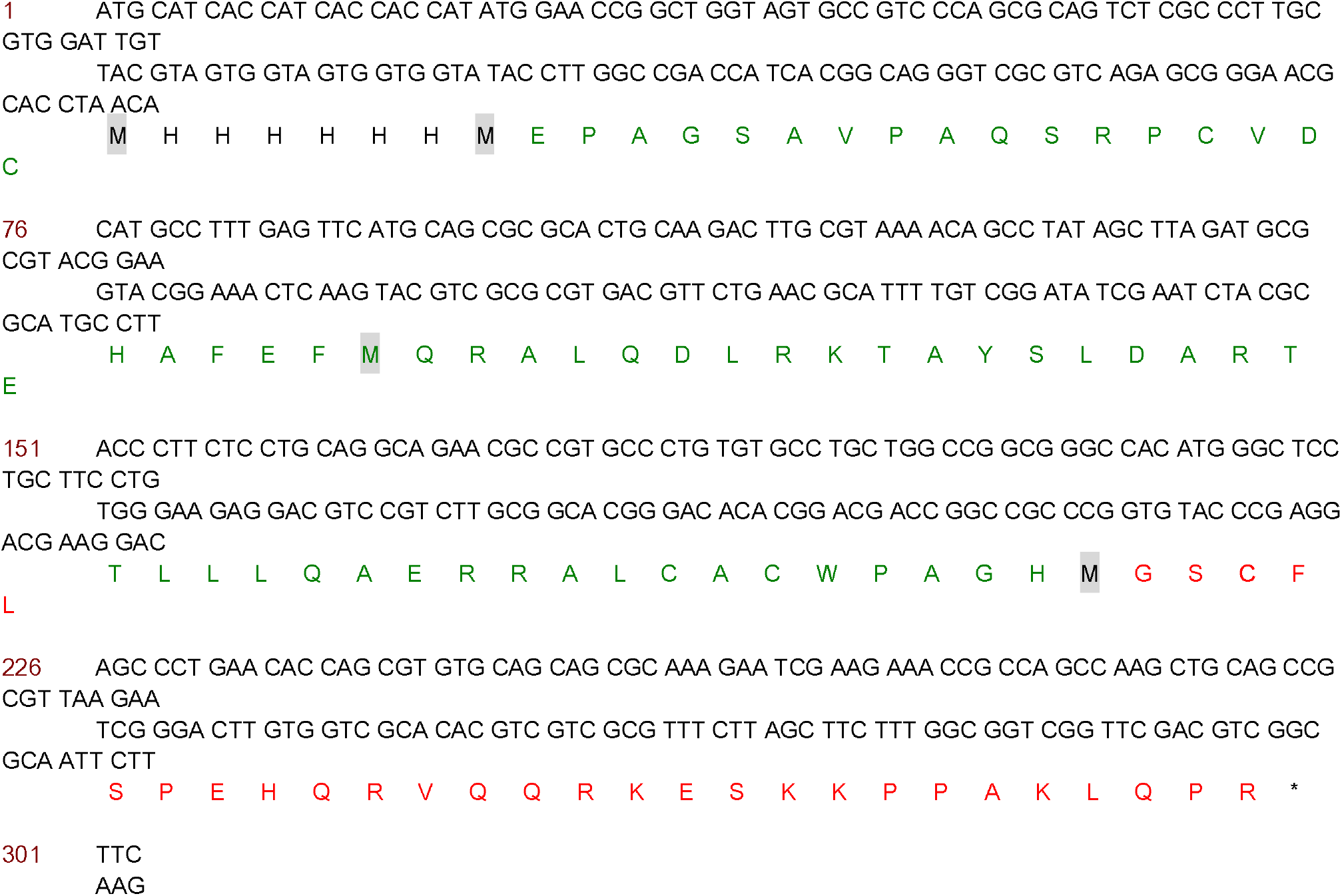
The nucleotide sequence and amino acid sequence of 6×His-C4ORF48-[S3C]UAG fusion protein overexpressed in *E. coli*. The amino acid sequence of [S3C]UAG is shown in red, and that of the predicted mature peptide of C4ORF48 is shown in green. Met residues for CNBr cleavage are shaded.

**Fig. S2.**
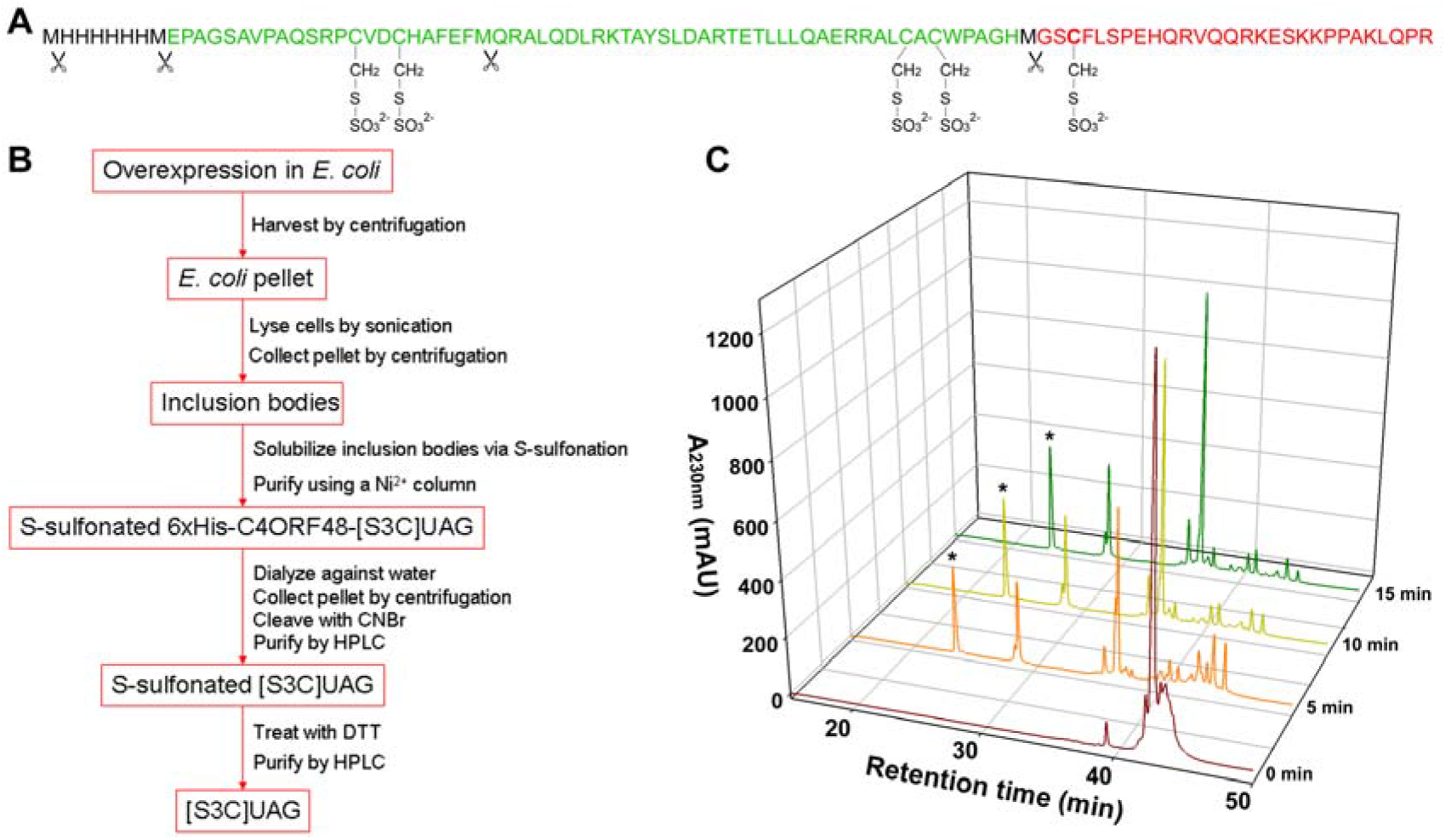
Preparation of [S3C]UAG by overexpression of a larger precursor in *E. coli* and subsequent CNBr cleavage. (**A**) Amino acid sequence of the *S*-sulfonated 6×His-C4ORF48-[S3C]UAG. The amino acids in the fusion protein are shown as one-letter code. The side-chain of Cys residues and the reversibly modified sulfonate moieties are shown. The CNBr cleavage sites are indicated by scissors symbols. (**B**) A flowchart for preparation of the mature [S3C]UAG by bacterial overexpression and subsequent chemical cleavage. (**C**) HPLC analysis of the reaction mixture after the *S*-sulfonated precursor was cleaved by CNBr for different times. The peak of the *S*-sulfonated [S3C]UAG is indicated by an asterisk.

## References

1 Kojima M, Hosoda H, Date Y, Nakazato M, Matsuo H, Kangawa K. (1999) Ghrelin is a growth-hormone-releasing acylated peptide from stomach. Nature 402: 656−660.

2 Howard AD, Feighner SD, Cully DF, Arena JP, Liberator PA, Rosenblum CI, Hamelin M, Hreniuk DL, Palyha OC, Anderson J, Paress PS, Diaz C, Chou M, Liu KK, McKee KK, Pong SS, Chaung LY, Elbrecht A, Dashkevicz M, Heavens R, Rigby M, Sirinathsinghji DJ, Dean DC, Melillo DG, Patchett AA, Nargund R, Griffin PR, DeMartino JA, Gupta SK, Schaeffer JM, Smith RG, Van der Ploeg LH. (1996) A receptor in pituitary and hypothalamus that functions in growth hormone release. Science 273: 974−977.

3 Yang J, Brown MS, Liang G, Grishin NV, Goldstein JL. (2008) Identification of the acyltransferase that octanoylates ghrelin, an appetite-stimulating peptide hormone. Cell 132: 387–396.

4 Gutierrez JA, Solenberg PJ, Perkins DR, Willency JA, Knierman MD, Jin Z, Witcher DR, Luo S, Onyia JE, Hale JE. (2008) Ghrelin octanoylation mediated by an orphan lipid transferase. Proc Natl Acad Sci USA 105: 6320–6325.

5 Ge X, Yang H, Bednarek MA, Galon-Tilleman H, Chen P, Chen M, Lichtman JS, Wang Y, Dalmas O, Yin Y, Tian H, Jermutus L, Grimsby J, Rondinone CM, Konkar A, Kaplan DD. (2018) LEAP2 Is an Endogenous Antagonist of the Ghrelin Receptor. Cell Metab 27: 461–469.

6 Wang JH, Li HZ, Shao XX, Nie WH, Liu YL, Xu ZG, Guo ZY. (2019) Identifying the binding mechanism of LEAP2 to receptor GHSR1a. FEBS J 286: 1332–1345.

7 M’Kadmi C, Cabral A, Barrile F, Giribaldi J, Cantel S, Damian M, Mary S, Denoyelle S, Dutertre S, Péraldi-Roux S, Neasta J, Oiry C, Banères JL, Marie J, Perello M, Fehrentz JA. (2019) N-Terminal Liver-Expressed Antimicrobial Peptide 2 (LEAP2) Region Exhibits Inverse Agonist Activity toward the Ghrelin Receptor. J Med Chem 62: 965–973.

8 Mani BK, Puzziferri N, He Z, Rodriguez JA, Osborne-Lawrence S, Metzger NP, Chhina N, Gaylinn B, Thorner MO, Thomas EL, Bell JD, Williams KW, Goldstone AP, Zigman JM. (2019) LEAP2 changes with body mass and food intake in humans and mice. J Clin Invest 129: 3909–3923.

9 Cornejo MP, Castrogiovanni D, Schiöth HB, Reynaldo M, Marie J, Fehrentz JA, Perello M. (2019) Growth hormone secretagogue receptor signalling affects high-fat intake independently of plasma levels of ghrelin and LEAP2, in a 4-day binge eating model. J Neuroendocrinol 31: e12785.

10 Islam MN, Mita Y, Maruyama K, Tanida R, Zhang W, Sakoda H, Nakazato M. (2020) Liver-expressed antimicrobial peptide 2 antagonizes the effect of ghrelin in rodents. J Endocrinol 244: 13–23.

11 Li HZ, Shou LL, Shao XX, Li N, Liu YL, Xu ZG, Guo ZY. (2021) LEAP2 has antagonized the ghrelin receptor GHSR1a since its emergence in ancient fish. Amino Acids 53: 939–949.

12 Abizaid A, Hougland JL. (2020) Ghrelin Signaling: GOAT and GHS-R1a Take a LEAP in Complexity. Trends Endocrinol Metab 31: 107–117.

13 Al-Massadi O, Müller T, Tschöp M, Diéguez C, Nogueiras R. (2018) Ghrelin and LEAP-2: Rivals in Energy Metabolism. Trends Pharmacol Sci 39: 685–694.

14 Cervone DT, Lovell AJ, Dyck DJ. (2020) Regulation of adipose tissue and skeletal muscle substrate metabolism by the stomach-derived hormone, ghrelin. Curr Opin Pharmacol 52: 25–32.

15 Yanagi S, Sato T, Kangawa K, Nakazato M. (2018) The Homeostatic Force of Ghrelin. Cell Metab 27: 786–804.

16 Delhanty PJ, Neggers SJ, van der Lely AJ. (2013) Des-acyl ghrelin: a metabolically active peptide. Endocr Dev 25: 112–121.

17 Delhanty PJ, Neggers SJ, van der Lely AJ. (2012) Mechanisms in endocrinology: Ghrelin: the differences between acyl- and des-acyl ghrelin. Eur J Endocrinol 167: 601–608.

18 Gortan Cappellari G, Barazzoni R. (2019) Ghrelin forms in the modulation of energy balance and metabolism. Eat Weight Disord 24: 997–1013.

19 Chen VP, Gao Y, Geng L, Parks RJ, Pang YP, Brimijoin S. (2015) Plasma butyrylcholinesterase regulates ghrelin to control aggression. Proc Natl Acad Sci USA 112: 2251–2256.

20 Eubanks LM, Stowe GN, De Lamo Marin S, Mayorov AV, Hixon MS, Janda KD. (2011) Identification of alpha2 macroglobulin as a major serum ghrelin esterase. Angew Chem Int Ed Engl 50: 10699–10702.

21 Satou M, Nishi Y, Yoh J, Hattori Y, Sugimoto H. (2010) Identification and characterization of acyl-protein thioesterase 1/lysophospholipase I as a ghrelin deacylation/lysophospholipid hydrolyzing enzyme in fetal bovine serum and conditioned medium. Endocrinology 151: 4765–4775.

22 De Vriese C, Gregoire F, Lema-Kisoka R, Waelbroeck M, Robberecht P, Delporte C. (2004) Ghrelin degradation by serum and tissue homogenates: identification of the cleavage sites. Endocrinology 145: 4997–5005.

23 Matsumoto M, Hosoda H, Kitajima Y, Morozumi N, Minamitake Y, Tanaka S, Matsuo H, Kojima M, Hayashi Y, Kangawa K. (2001) Structure-activity relationship of ghrelin: pharmacological study of ghrelin peptides. Biochem Biophys Res Commun 287: 142–146.

24 Ahangarpour M, Kavianinia I, Harris PWR, Brimble MA. (2021) Photo-induced radical thiol-ene chemistry: a versatile toolbox for peptide-based drug design. Chem Soc Rev 50: 898–944.

25 Nolan MD, Scanlan EM. (2020) Applications of Thiol-Ene Chemistry for Peptide Science. Front Chem 8: 583272.

26 Kowalczyk R, Harris PWR, Williams GM, Yang SH, Brimble MA. (2017) Peptide Lipidation - A Synthetic Strategy to Afford Peptide Based Therapeutics. Adv Exp Med Biol 1030: 185–227.

27 Hermant YO, Cameron AJ, Harris PWR, Brimble MA. (2020) Synthesis of Antimicrobial Lipopeptides Using the “CLipPA” Thiol-Ene Reaction. Methods Mol Biol 2103: 263–274.

28 Yim V, Hermant YO, Harris PWR, Brimble MA. (2021) Synthesis of Lipopeptides by CLipPA Chemistry. Methods Mol Biol 2355: 253–263.

29 Li HZ, Shou LL, Shao XX, Liu YL, Xu ZG, Guo ZY. (2020) Identifying key residues and key interactions for the binding of LEAP2 to receptor GHSR1a. Biochem J 477: 3199–3217.

30 Liu Y, Shao XX, Zhang L, Song G, Liu YL, Xu ZG, Guo ZY. (2015) Novel bioluminescent receptor-binding assays for peptide hormones: using ghrelin as a model. Amino Acids 47: 2237–2243.

31 Li HZ, Shao XX, Shou LL, Li N, Liu YL, Xu ZG, Guo ZY. (2021) Unusual orthologs shed new light on the binding mechanism of ghrelin to its receptor GHSR1a. Arch Biochem Biophys 704: 108872.

32 Matthews RP, Guthrie CR, Wailes LM, Zhao X, Means AR, McKnight GS (1994) Calcium/calmodulin-dependent protein kinase types II and IV differentially regulate CREB-dependent gene expression. Mol Cell Biol 14: 6107–6116.

33 Holst B, Brandt E, Bach A, Heding A, Schwartz TW. (2005) Nonpeptide and peptide growth hormone secretagogues act both as ghrelin receptor agonist and as positive or negative allosteric modulators of ghrelin signaling. Mol Endocrinol 19: 2400–2411.

